# Syntaxin-3 regulates Tight Junction Assembly in Human Retinal Pigment Epithelium

**DOI:** 10.64898/2026.01.30.702706

**Authors:** Athira Ramesh, Suganya Sivagurunathan, Navya Valavath Baburajan, Subbulakshmi Chidambaram

## Abstract

The retinal pigment epithelium (RPE) consists of polarized epithelial cells, serving as a support system for photoreceptor maintenance, where the polarity is contributed by the distribution of syntaxins (STX) within the cell. STX3, a known regulator of apical trafficking in epithelial cells, was previously understood to be absent in human RPE cells, with its functions thought to be compensated by STX1A. However, our results on SNARE mRNA expression profile in RPE detected the presence of 2 splice variants of STX3. Further investigation in donor retina, primary hRPE, and ARPE-19 cells revealed detectable levels of STX3 mRNA and protein. STX3 knockdown in ARPE19 resulted in a significant reduction of tight junction (TJ) proteins, compromising TJ assembly, highlighting the critical role of STX3 in maintaining RPE integrity. In addition, immunoprecipitation followed by LC-MS/MS analysis revealed that STX3 and STX1A have a distinct novel protein interactome in RPE. This study identified unique and shared interactants for STX3 and STX1A, suggesting a broader role for RPE beyond its traditional photoreceptor support function. This further emphasises the biological significance of STX1A and STX3 in maintaining retinal homeostasis, which could facilitate the development of novel therapeutic strategies for retinal disorders.

**Significance:** This study identified the presence of STX3 in the human RPE cells, which was previously reported to be absent. Further, we demonstrated that STX3 knockdown in ARPE19 cells disrupted TJ assembly, highlighting its potential role in preserving RPE cell polarity and structural integrity, challenging the notion that STX3 functions were thought to be compensated by STX1A. Moreover, immunoprecipitation followed by LC-MS/MS analysis in RPE identified the protein interaction networks of both STX1A and STX3. Interestingly, unique and shared interactants, including proteins associated with neuronal plasticity, indicated unidentified functions of STX3 and STX1A in RPE. This suggests that they might perform both overlapping and distinct functions for maintaining RPE cell integrity and thus retinal homeostasis. Overall, our preliminary findings challenge the established view that STX3 is non-existent in RPE cells and initiate new directions for exploring the multifaceted and potentially non-redundant functions of STX3 in RPE.

## Introduction

The retinal pigment epithelium (RPE) is a layer of polarized epithelial cells located between the choroidal capillaries and the photoreceptors. The distinctive structural polarity and specialized trafficking mechanisms exhibited by RPE are crucial for maintaining retinal homeostasis, health, and integrity [1,2], distinguishing it from most other extraocular epithelial cells [3]. The major function of the RPE is to support photoreceptors by providing nourishment, recycling retinoids, and facilitating the phagocytosis and renewal of photoreceptor outer segments (POS), all of which are essential for maintaining photoreceptor functioning [2]. Phagocytic clearance of approximately 10% of photoreceptor outer segment debris by RPE is facilitated by the polarised expression of proteins concentrated at its apical membrane [4]. Moreover, the tight junctions (TJs) also play a significant role in these transport events, which are located on the apical-lateral sides of the RPE. This divides the RPE into apical and basolateral domains by compartmentalizing receptors and ion channels into two distinct parts, thereby facilitating the polarised secretion of reinforcement factors, such as VEGF and PEDF. Interestingly, RPE exhibits ‘reversed polarity’, with proteins like Na/K-ATPase, N-CAM, EMMPRIN, etc., typically expressed in the basolateral membrane in other epithelial cells, are localised on the apical membrane of RPE. Notably, the reverse polarity of ion channels such as Na/K-ATPase in RPE provides an accurate ionic environment and appropriate niche to support phototransduction, thereby playing an important role in cargo transport and maintaining RPE homeostasis [5-7].

In addition, trafficking proteins such as SNAREs, including syntaxins (STXs), VAMPs, SNAPs, and RABs, also contribute to the distinctive phenotypes and polar organisation of epithelial cells, including RPE and neurons, which are implicated in specialised, polarised trafficking and are pivotal for their functioning. Among these, STXs have a polarised distribution in epithelial cells. STX1A, a Qa SNARE, which is normally restricted to neurons and neuroendocrine cells, is expressed in the RPE apical membrane, and both in inner and outer plexiform layers of the retina [9,10,13,42]. In neurons, it is involved in synaptic transmission, along with SNAP25 and VAMPs [14,15]. Studies in MDCK cells on STXs have stated that STX4 is involved in TGN to basolateral transport [16]. STX3, an apical targeting protein, facilitates transport from the TGN to the apical membrane [17] and is crucial for preserving both epithelial polarity [12] and neuronal cell polarity with associated trafficking processes [8]. Furthermore, STX2 and STX4 are observed to localise near the TJ of RPE, implicating the possibility of vesicle fusion or preferred sites for exocytosis in the proximity of TJs [10]. Additionally, STX3 plays an extensive role in establishing polarised skin epithelial architecture and maintaining TJ barrier integrity [18], and STX2, expressed in the basolateral domain of RPE, exhibited reverse polarity based on epithelial cell type [19, 10]. Previous research has identified STX1A, STX1B, STX2, and STX4 in RPE cells, where they contribute to cell polarity and trafficking. Notably, a recent study highlighted the critical role of STX3 in the proper delivery of disc proteins to the outer segments of photoreceptors [20]. Despite this, the detailed mechanisms governing these trafficking pathways remain only partially understood. Furthermore, a study by Perez-Hurtado et al. revealed that a single functional copy of STX3 is adequate to sustain the survival of mouse photoreceptors [21].

Intriguingly, Low et al. conducted a comparative analysis of the expression and distribution of SNAREs in the rat RPE cell line (RPE-J) and kidney epithelial cells (MDCK), exposing significant differences in SNARE expression and localisation. These findings underscore the distinct characteristics and specialised polarity of RPE cells compared to other epithelial cell types. However, the expression profile of STXs in RPE-J revealed the presence of STX1A and STX1B, while STX3 was not detected. Consequently, the study concluded that the absence of both mRNA and protein presence of STX3 in RPE-J cells suggests that the STX3-dependent pathways are likely absent in RPE cells [10]. Interestingly, a few studies have suggested that STX1A, which is structurally similar to STX3 and which forms a functional SNARE complex with SNAP25 and VAMP2 [22-26], could compensate for the latter’s function in RPE [10]. However, the broader functions of STX3 and STX1A in RPE cells and their overall significance in retinal physiology warrant further investigation. Interestingly, our experiments in primary RPE cells, donor RPE tissue, and the ARPE19 demonstrated the presence of STX3 at both transcript and protein levels, and its knockdown had a significant effect on TJ assembly. Therefore, we propose that STX3 maintains unique and specific functions within RPE cells that are not compensated by other proteins, which could influence the RPE cellular integrity and retinal physiology. In addition to this, exploring the STX3 and STX1A proteome in RPE cells is essential to comprehend their significance in retinal physiology and would facilitate the development of novel therapeutic options for retinal disorders.

## Materials & Methods

### Primary hRPE culture

Donor eye tissue was procured from CU Shah Eye Bank, Sankara Nethralaya, Chennai, India, under the Ethics subcommittee (Institute Review Board), Vision Research Foundation. For primary human RPE (hRPE) isolation, we collected eye tissue from three donors: two males (ages 32 and 35) and one female (age 42). The eyes were obtained within 6 hours after postmortem, and the processing was initiated within 2 hours of collection. The explant culture technique described by Zhu et al [27] was adapted for a cell isolation method, with a few modifications [28]. In brief, to preserve the delicate layer of RPE, the front portions of the eye, vitreous, and retina were gently removed. After thoroughly washing the remaining eye globe with antibiotic-enriched PBS solution, the eye was cut into flower-like sections. Each piece was placed face-down on six-well plates that had been precoated with fibronectin (10μg/ml in PBS). Further, the tough outer sclera layer was delicately peeled away, leaving the fragile RPE layer intact. To facilitate the cell growth, 100μl of endothelial growth medium supplemented with growth factors was added, and the cultures were undisturbed for 4-5 hours. Later, 1ml of medium was added, and after allowing the explants to attach overnight, the tissue pieces were removed. For continued growth, the emerging cells were transferred to flasks coated with 0.1% gelatin and nourished with DMEM-F12 medium (Sigma-Aldrich, USA) supplemented with 14.2 mM sodium bicarbonate, 10% BSA, and antibiotics was used to grow hRPE cells isolated from donor RPE tissue and ARPE19 cells (ATCC-CRL-2302) were cultured to confluence. hRPE cells were used up to 6 passages for the study.

### qRT-PCR and Western blot analysis

SYBR green (Bio-Rad, USA) assays were set up using the cDNA with the primers listed in the Sup. Table 1. The fold change of the transcripts in qPCR was calculated based on the relative quantification (2-ΔΔCq) method in qPCR with the internal control *18S rRNA*. For Western blot analysis, 50μg of protein was used per assay. The blots were probed against anti-rabbit *STX3* and *STX1A* antibodies (a generous gift from Prof. Reinhard Jahn, MPIBPC, Göttingen), anti-mouse *β-Actin* antibody (SantaCruz Biotechnology, 1:1000), and anti-rabbit and anti-mouse secondary antibodies (SantaCruz Biotechnology, 1:10,000) were used for detection.

### Immunofluorescence

RNAi studies were performed using *si-STX3* (siRNA target sequence-AAGGGCCAACAACGTCCGGAA) [29] at a 5 nM concentration. ARPE19 cells were grown on coverslips until reaching 75% confluence, then transfected with either *si-STX3* or *si-Scrambled*. 48 hours post-transfection, cells were fixed with 4% paraformaldehyde and permeabilized with 0.2% Triton-X 100. Subsequently, cells were incubated overnight with primary antibodies, anti-rabbit anti-*ZO1*, and anti-*Occludin* (Invitrogen, 1:100 dilution), followed by incubation with FITC-conjugated secondary antibodies (1:400 dilution, Invitrogen, USA) for 2 hours at room temperature. DAPI staining was performed before mounting the coverslips onto glass slides with ProLong anti-fade reagent (Invitrogen, USA). Imaging was conducted at 63X magnification using an LSM710 confocal laser scanning microscope (Carl Zeiss, Germany).

### Immunoprecipitation and LC-MS/MS

Immunoprecipitation of STX3 and STX1A was performed using ARPE19 cells. The ARPE19 cell lysate was prepared in RIPA buffer and incubated on ice for 20 minutes. The lysate was then centrifuged at 14,000 rpm for 20 minutes at 4°C to remove the debris. For comparison, 1/10th of the protein fraction was saved as Input. The remaining protein fraction was separated into three aliquots, each incubated separately with 2μg of anti-*STX1A*, 2μg of anti-*STX3* antibodies, and 1.5μl of Rabbit IgG Isotype control. The mixture was incubated at 4°C for 2 hours on a rotating apparatus. Following this, 30μl of Protein A/G Plus Agarose beads (Santa Cruz Biotechnology) were added, and the mixture was incubated overnight at 4°C. The following day, the beads were washed five times with 1x RIPA buffer containing 1x protease inhibitor. The enriched proteins were then combined with loading dye, heated for 5 minutes at 94°C, and used for SDS-PAGE gel for analysis. A Coomassie blue-stained SDS-PAGE gel of the immunoprecipitated samples has been included in the sup. Fig.3.

The samples were prepared for in-gel digestion and subsequently analyzed by LC-MS/MS. The peptides were analysed using a Q-Exactive Plus hybrid quadrupole-Orbitrap mass spectrometer, which was coupled with a Nano-LC pump from EASY-nLC, and manufactured by Thermo Fisher Scientific in Germany. Full-scan MS spectra were acquired in the range of 350–1600 m/z at a resolution of 70,000. The top 20 (in-gel digested samples) most intense peaks from the survey scan were selected for fragmentation with Higher-energy Collisional Dissociation with a normalized collision energy of 28% and isolation window 1.6 m/z. The dynamic exclusion time for precursor ions selected for MS/MS fragmentation was 30s. Automatic gain control target value and maximum ion injection time for MS and MS/MS were 1 × 10^6^ and 1 × 10^4^, respectively. Proteomic data were analyzed using FunRich_3.1.3 [30] for functional enrichment and network analysis.

### Homology modelling and molecular docking

Modeller 9.19 (https://salilab.org/modeller/9.19/release.html#windows) [31,32] and IntFOLD (http://www.reading.ac.uk/bioinf/IntFOLD/) [33] servers were employed for homology modelling and validated through RAMPAGE [34] and ProSA [35,36]. Protein sequences and templates for homology modelling were retrieved from NCBI and Protein Data Base (PDB), respectively. The disorder in the STX3 protein was predicted by DisEMBL [37]. The ‘easy interface’ of the HADDOCK web server (https://rascar.science.uu.nl/auth/register/haddock) [38] and LZerD (https://lzerd.kiharalab.org/about/) [39] webservers was used to execute protein-protein interactions involving STX3 (PDBid:1ez3, 1s94 as template) against Dynamin (DNM1) (PDBid:5d3q as positive control), CHM4B (PDBid:4abm, 3UM3), Coronin2B (CORO2B) (PDBid:2aq5, 2b4e), and KIDINS220 (IntFOLD server because of structure complexity), STX1A with SERPING1 (P05155), SLC1A3 (P43003) and PKN3 (Q6P5Z2) and both STX1A and STX3 with the common interactants KDELR2 (P33947), AMBP (P02760), LRG1 (P02750) and NSF (P46459). Docked structures were selected based on the HADDOCK score, which aggregates various interaction energies and then analyzed using PDBsum (https://www.ebi.ac.uk/thornton-srv/databases/pdbsum/) [40] and PDBePISA (https://www.ebi.ac.uk/pdbe/pisa/) [41]. No molecular dynamics simulation studies were done for the protein structures.

## Results

### Expression profile of SNARE mRNAs and validation of STX3 expression in RPE

Given the critical role of SNARE proteins in regulating epithelial cell polarity and vesicular trafficking, we analysed the expression of SNARE genes in RPE cells using qRT-PCR. Expression of Q_a_ SNAREs (STX1A, STX2, STX3 variants 1 and 2, STX4, STX7, STX8, STX12, STX16), Q_b_ SNAREs (SNAP23, SNAP25, SNAP29), Q_c_ SNARE STX6, R-SNAREs (VAMP1, VAMP2, VAMP3, VAMP7, VAMP8), and RABs (RAB1, RAB3A, RAB5, RAB6, RAB8, RAB11) was detected in ARPE-19 cells (Fig.1a). Intriguingly, STX3 was not detected in rat RPE-J cells in a previous study [10], and its presence in the human RPE has not been experimentally validated until now. Notably, our data displayed the expression of two variants (among 4 existing variants) of *STX3* transcript in ARPE19 cell line, primary hRPE cells, and donor RPE tissue, as demonstrated by RT-PCR (Fig. 1b), which was further validated by qPCR (Fig.1c). Additionally, we performed Western blot analysis and confirmed the presence of STX3 protein in ARPE19, hRPE and donor RPE tissue (Fig.1d). Thus, our results demonstrated the presence of STX3 in human RPE.

**Fig. 1.**
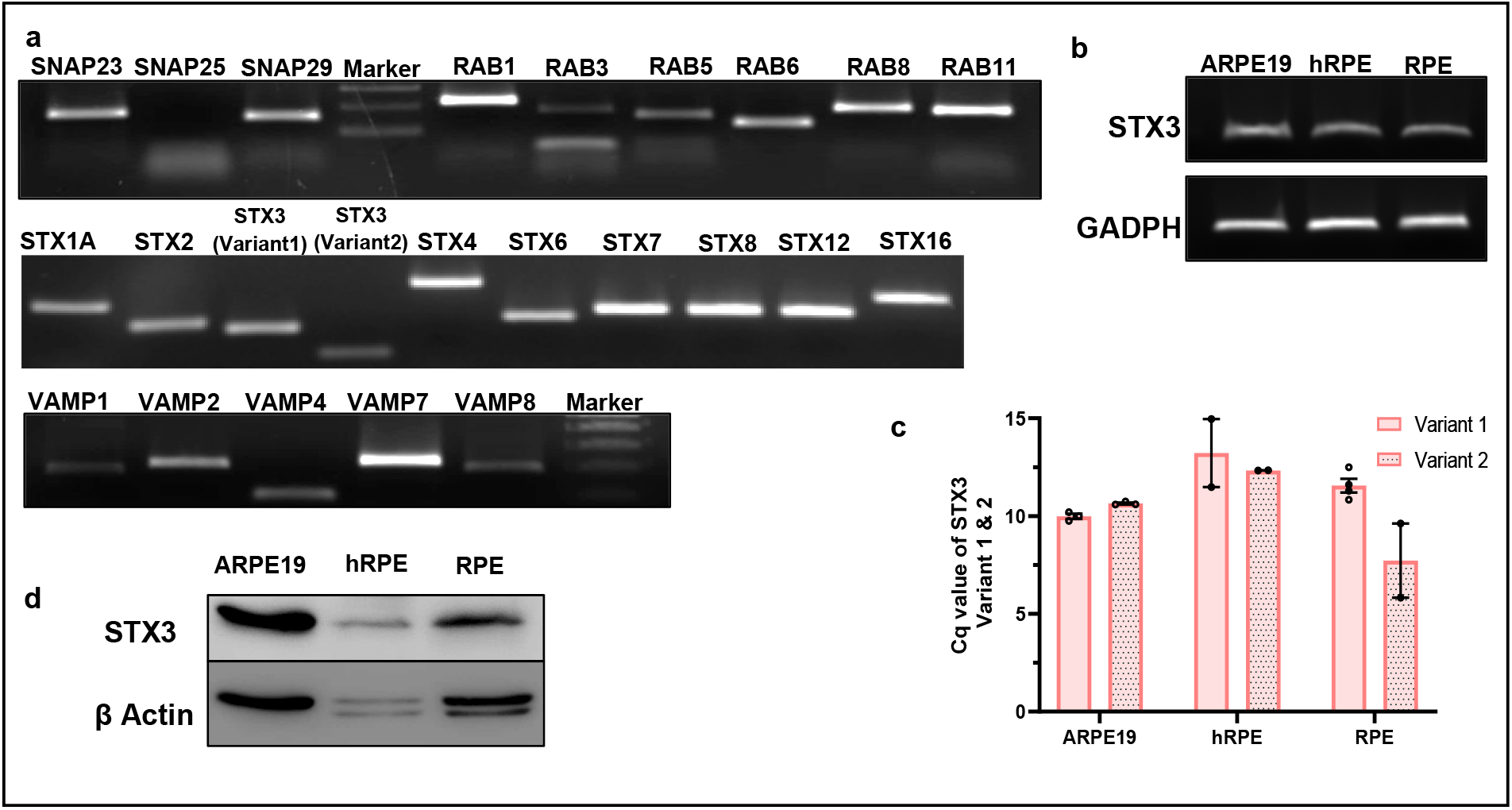
Identification and validation of STX3 expression in RPE. (**a)** The RT-PCR results depict the expression of various SNARE proteins, SNAP 23,25 and 29, Rabs 1, 3, 5, 6, 8 and 11, Syntaxins 1A, 2, 3 (variant1 and 2), 4, 6, 7, 8, 12 and 16, VAMPs 1, 2, 3, 7 and 8 and STX17 in ARPE19 cell line. (**b)** RT-PCR of STX3 in ARPE19, primary hRPE cells, and donor RPE tissue. (**c)** qPCR analysis of STX3 variant 1 and 2 in ARPE19, primary hRPE cells, and donor RPE tissue. The Cq values were normalized with 18S rRNA as the internal control, ARPE19, n=3, hRPE, RPE layer n=2. (**d)** STX3 protein expression was further validated by the Western blot in ARPE19, primary hRPE cells, and donor RPE tissue

### STX3 silencing affects the tight junction integrity in ARPE19

Previous studies have indicated multiple roles of STX3 in various cell types, including its involvement in polarity [12], apical trafficking [3,42]. and barrier formation [18]. We anticipated that STX3 would exhibit similar functions in RPE, maintaining cell integrity by regulating the abundance of TJ proteins and their assembly. To investigate this, we silenced STX3 with siRNA (Fig. 2a,b) and examined the presence and assembly of TJ proteins through immunofluorescence. Interestingly, the confocal images revealed reduced immunostaining at cell-cell contact sites, attributed to the noticeable depletion of the TJ proteins, ZO1, and occludin upon *STX3* silencing (Fig. 2c). This significant reduction could destabilize the integrity of TJ assembly in ARPE19, indicating that STX3 is crucial for maintaining the cellular integrity of RPE which might affect the polarity of the cell. Thus, our data indicate that STX3 could be important for maintaining the cellular integrity of RPE.

**Fig. 2.**
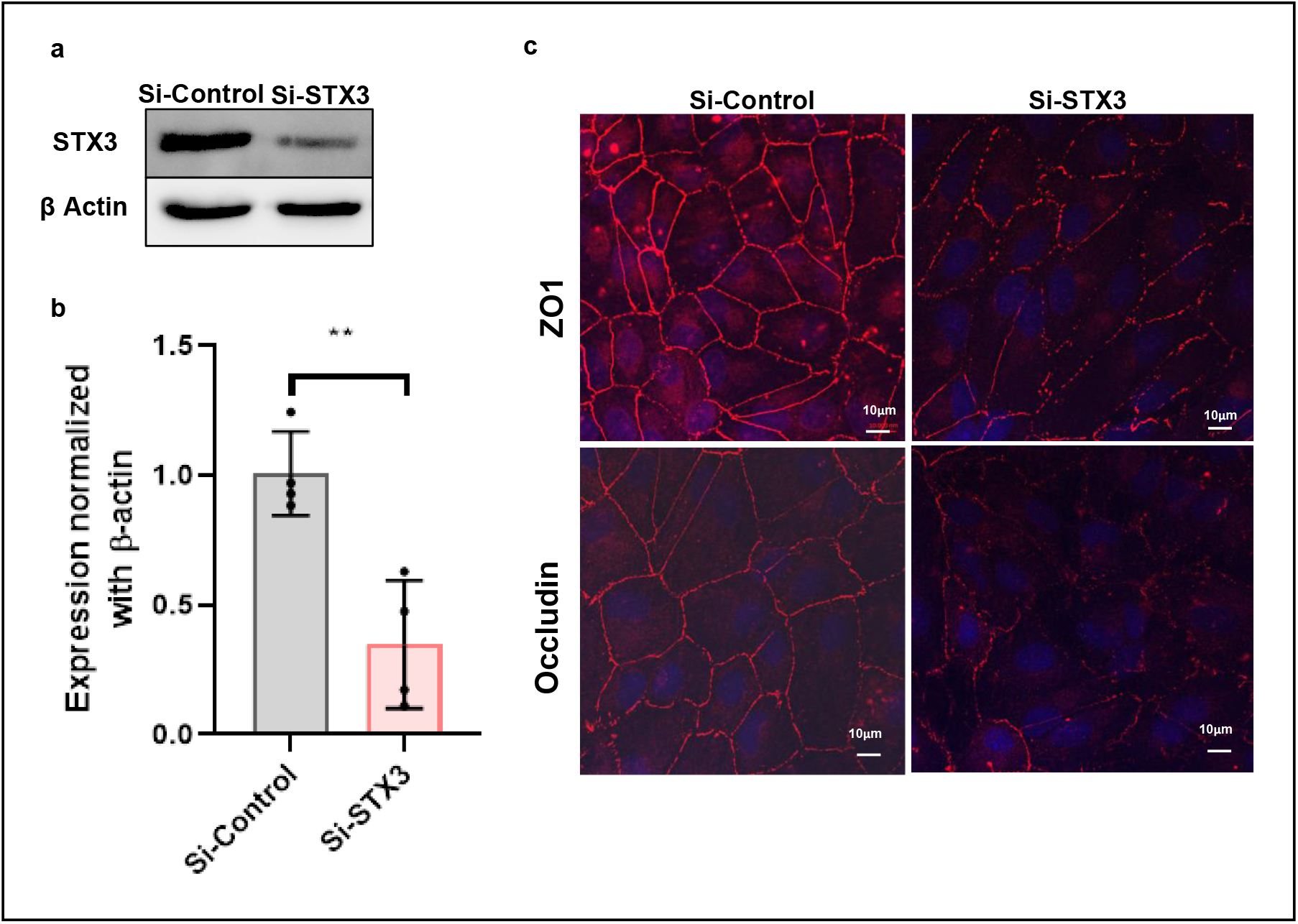
STX3 silencing affects the tight junction integrity in ARPE19. (**a)** The Western blot analysis showing siRNA-mediated knockdown of STX3 in ARPE19 cells. (**b)** Quantification for the Western blot analysis was done using ImageJ software, normalised with loading control *β-Actin*, and the graph was plotted with GraphPad Prism 8.4.2. Statistical significance for siRNA-mediated knockdown of STX3 in ARPE19 cells was calculated by T-test with P<0.004, with N=4, biological replicates. (**c)** Representative confocal images of confluent ARPE19 cells stained for TJ proteins, ZO1 and Occludin in control and *siRNA-*mediated STX3 knockdown. The nuclear staining was accomplished using DAPI. Expression of ZO1 and Occludin was observed to be reduced in *si-STX3* compared to *si-Control*.

### Identification and comparison of interacting proteins of STX3 and STX1A in ARPE19 by immunoprecipitation and LC-MS/MS

To further demarcate the specific functions of STX3, we conducted immunoprecipitation in ARPE19 for both STX3 (Fig. 3a) and STX1A (Fig. 3b), followed by LC-MS/MS. To identify and compare the unique protein interaction partners of STX3 and STX1A, we set a peptide count threshold of three for the LC-MS/MS data analysis. This analysis revealed 106 proteins uniquely interacting with STX3, 30 with STX1A, and 33 shared between both STX family proteins (Fig. 3c). The top novel interacting proteins identified were sorted by the highest number of matching peptides and listed in Table 1. DNM1, an already reported interactant of STXs, was also identified along with the other proteins [43,63]. Notably, a higher proportion of STX3 interactors exhibited neuronal origins, suggesting potential novel functions for STX3 in RPE. Functional enrichment analysis using FunRich revealed the involvement of STX3 and STX1A in vesicular transport, cell communication, cell growth, and neurotransmitter transport (Fig. 3d, e, f).

**Fig. 3.**
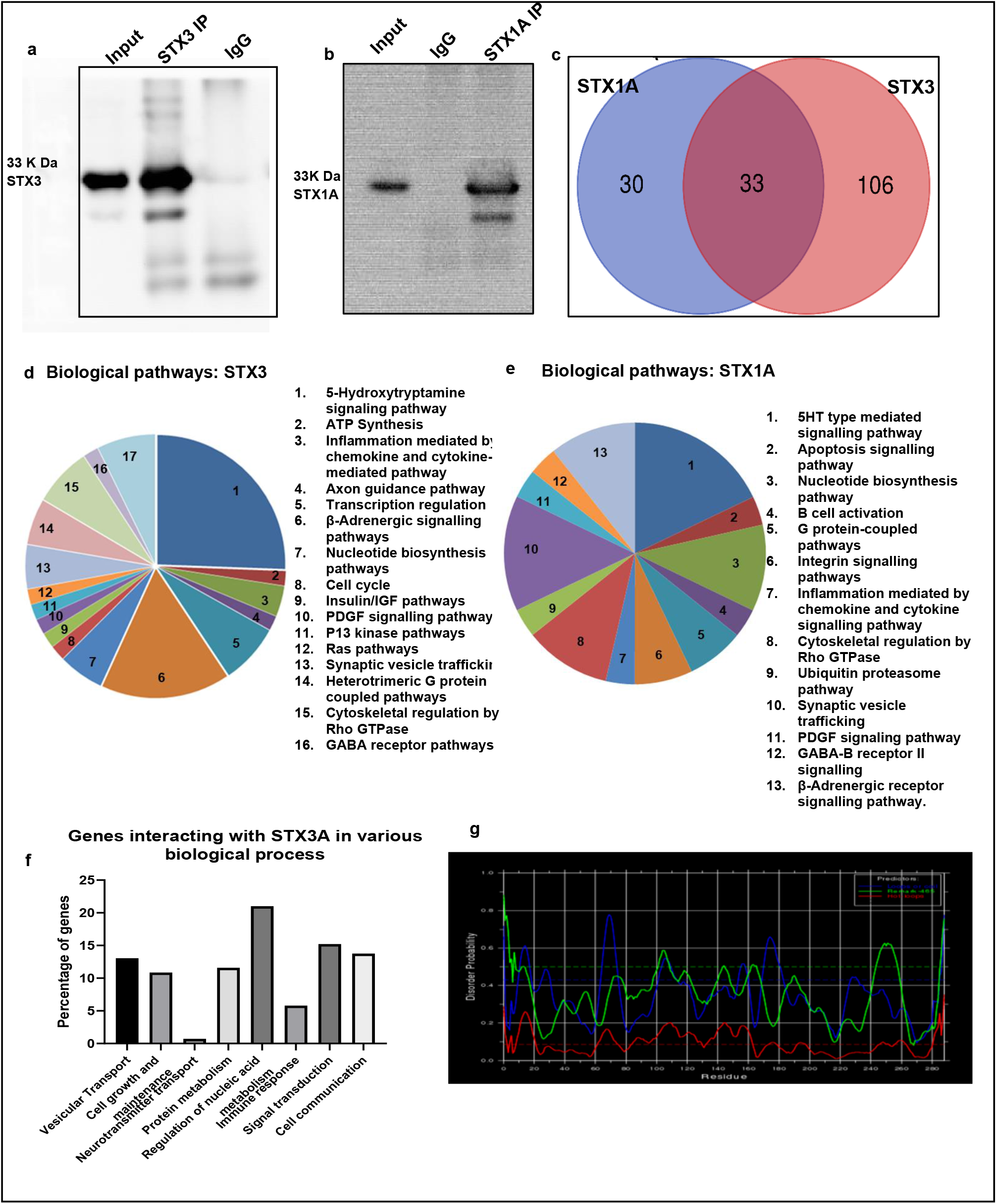
Immunoprecipitation and proteomics of STX3 and STX1A in ARPE19. (**a)** The Western blot image of STX3 immunoprecipitation in ARPE19. (**b**) The Western blot image of STX1A immunoprecipitation in ARPE19. **(c)** Venn diagram showing the number of common and specific proteins interacting with STX3 and STX1A identified through immunoprecipitation followed by LC/MS/MS. Functional analysis of interacting proteins of STX3 and STX1A: Predicted biological pathways for STX3 **(d)** and STX1A **(e)** using FunRich. (**f)** Percentage of genes interacting with STX3 in various biological processes. **(g)** The disordered region in the STX3 structure was predicted by DisEMBL with disorder probability on the x-axis and residues on the y-axis. The regions above 0.5 disorder probability are considered as intrinsically disordered regions. Predictors: the blue line - loops or coils, the green line- Remark-465, the red line-hot loops.

**Table. 1:**
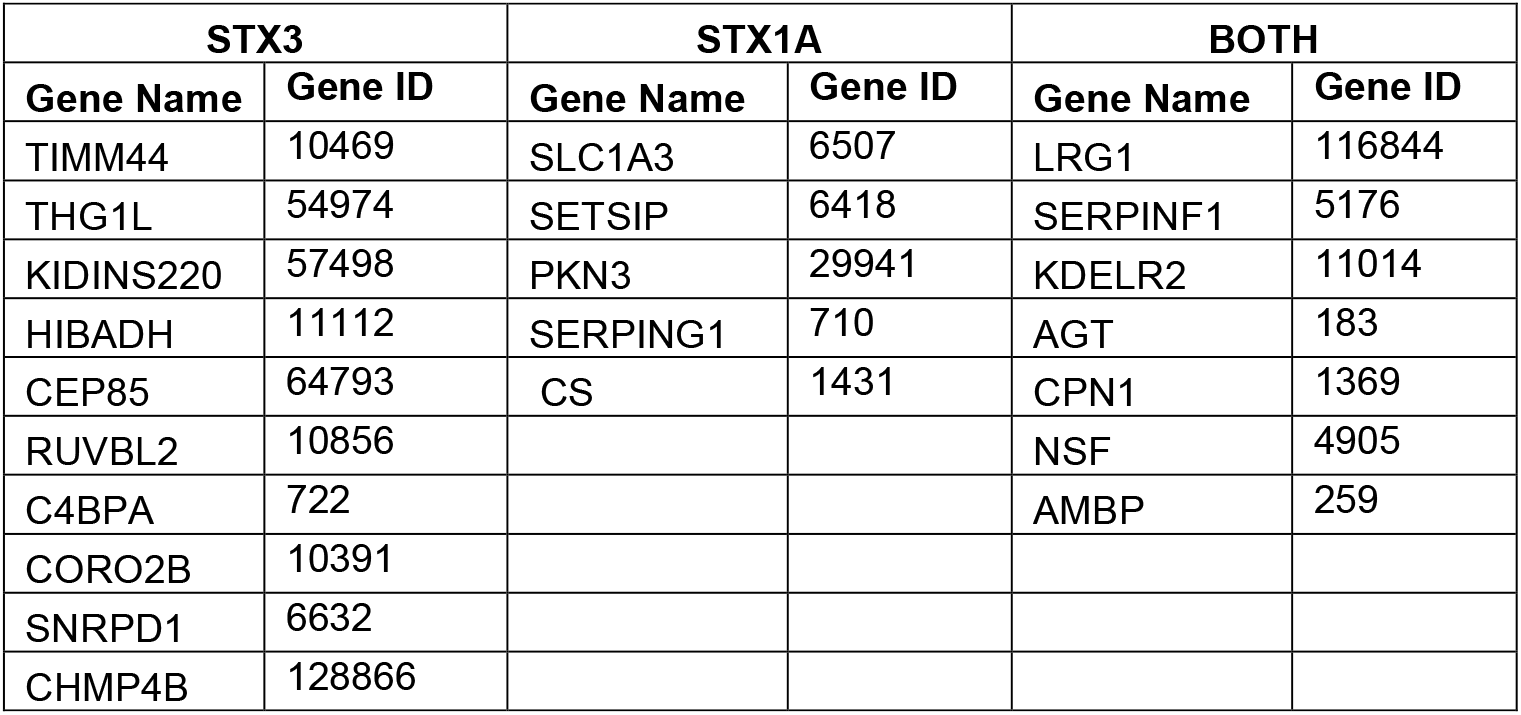
List of interacting protein partners of STX3 and STX1A identified through LC-MS/MS.

| STX3 |  | STX1A |  | BOTH |  |
| --- | --- | --- | --- | --- | --- |
| Gene Name | Gene ID | Gene Name | Gene ID | Gene Name | Gene ID |
| TIMM44 | 10469 | SLC1A3 | 6507 | LRG1 | 116844 |
| THG1L | 54974 | SETSIP | 6418 | SERPINF1 | 5176 |
| KIDINS220 | 57498 | PKN3 | 29941 | KDELR2 | 11014 |
| HIBADH | 11112 | SERPING1 | 710 | AGT | 183 |
| CEP85 | 64793 | CS | 1431 | CPN1 | 1369 |
| RUVBL2 | 10856 |  |  | NSF | 4905 |
| C4BPA | 722 |  |  | AMBP | 259 |
| CORO2B | 10391 |  |  |  |  |
| SNRPD1 | 6632 |  |  |  |  |
| CHMP4B | 128866 |  |  |  |  |

Our proteomic analysis identified a surprising array of novel STX3 interactors with predominantly neuronal characteristics, including proteins implicated in neuronal survival, migration, development, and plasticity. From our study, we found that STX3 interacts with a cluster of proteins, including KIDINS220, CHMP4B, CORO2B, THG1L, NME1, and GNAO1, that display strong neuronal specificity and have been linked to maintaining neuronal plasticity. STX1A interacts with neuronal proteins involved in amino acid transport, such as SLC1A3, as well as cell adhesion protein PKN3, and serine protease inhibitors, such as SERPING1, etc. Both STX1A and STX3 interact with proteins essential for retinal homeostasis and Golgi-mediated trafficking, including AMBP, KDELR2, LRG1, and NSF (Table. 1, Sup. Data 1). Moreover, the FunRich analysis revealed that the STX3 interactome is enriched with purine and pyrimidine metabolism-induced signalling pathways, which are essential for neuronal plasticity [45,46]. To assess the feasibility of the interactions of STX3 and STX1A with their respective partners and to explore their potential neuronal-like functions in RPE, preliminary molecular docking studies were performed. CHMP4B, CORO2B, and KIDINS220 interact with STX3, SLC1A3, and PKN3, while SERPING1 interacts with STX1A and KDELR1. Additionally, LRG1, NSF, and AMBP interact with both protein groups. For reference, DNM1 and NSF, known STXs interactor [43,47,48,94], were included in the analysis as a positive control. The studies demonstrated that STX3 and STX1A interact favourably with their specific binding partners, with binding affinities comparable to their interaction with DNM1 (Sup. Table.3a,3b, Sup. Fig.4). The analysis showed that these interactions primarily could involve the N-terminal domain and SNARE motif of STX3. DisEMBL analysis predicted a high level of structural disorderness in the H_abc_ domain at the N-terminal of STX3, particularly in the residues 65-75, implying flexibility for interacting with diverse protein partners (Fig. 3g).

## Discussion

Retinal STX expression remains poorly characterised, and there are no prior reports documenting STX3 expression in the RPE or examining the functional significance of STX3 and STX1A in RPE cells and their potential impact on retinal physiology. While STX3 was previously reported to be undetectable in rat RPE-J cells, and STX1A was proposed as a potential compensatory factor, our current findings demonstrate the presence of both STX3 mRNA and protein in ARPE-19, primary hRPE cells, and donor RPE tissue. Further experiments indicate that STX3 could play a critical role in maintaining TJ integrity in RPE cells. In addition, we have identified novel interacting partners of both STX1A and STX3 in the RPE, highlighting new avenues of research into their functional role in retinal physiology and epithelial homeostasis. The discrepancy in STX3 expression in rat and human RPE may reflect the differences in the interspecies retinal development [10].

Humans and rodents exhibit fundamentally distinct patterns of retinal development. In humans, cone photoreceptors develop earlier than rods, and the central retina is cone-dominant. Additionally, the RPE exhibits uniform pigmentation. In contrast, rodent retinas are rod-dominant, with rods developing earlier than cones, and exhibit regional variations in RPE pigmentation. Furthermore, in humans, developmental genes exhibit prolonged expression windows, and most RPE maturation markers follow a temporally regulated expression pattern. In contrast, rodents undergo a more rapid developmental transition, with earlier appearance of RPE maturation markers [49-52]. Moreover, hRPE comprises heterogeneous cell populations exhibiting varied isoform expression of different proteins. These differences could account for the absence of STX3 expression in RPE-J cell lines. Functional studies using siRNA-mediated STX3 knockdown revealed its essential role in maintaining RPE barrier integrity by regulating ZO-1 and occludin expression and localisation. As tight junctions function as a physical barrier separating the apical and basolateral regions, they restrict solute movement across the epithelium. This regulation is vital for maintaining cellular polarity, which is crucial for the integrity and proper functioning of RPE. These results, along with recent findings highlighting involvement of STX3 in barrier formation in skin epithelium [18], strongly support the hypothesis that STX3 is essential for RPE barrier function, modulation, and polarity, thus maintaining retinal physiology. In addition to this, a few studies have suggested that STX1A, which is structurally similar to STX3, could compensate for the latter’s function in RPE [10,22]. However, our results demonstrate that silencing STX3 leads to a loss of TJ integrity, underscoring its critical functional role despite the presence of STX1A. Even though STX1A and STX3 have structural similarities, STX1A is mainly involved in synaptic vesicle fusion during neurotransmitter release, whereas STX3 localises to the apical plasma membrane of polarised cells. Both the STX1A and STX3 share common SNAP and VAMP partners in SNARE complex formation; however, the Munc proteins, which serve as both regulators and facilitators of SNARE-mediated membrane fusion, exhibit distinct binding specificities. Munc18-1 binds to STX1A, whereas Munc18-2 partners with STX3 [116-118]. A recent report has established that a rare mutation of STX3 in exon 9 causes a severe congenital diarrhoea called Microvillus inclusion disease, despite the presence of STX1A, due to the defects in the apical membrane trafficking in the epithelial cells [53]. Furthermore, the studies in neurons demonstrated that STX1A is essential in neurotransmission and synaptic plasticity, while STX3 is dispensable in hippocampal neurons [54-56]. These findings suggest that STX1A and STX3 could operate in complementary or overlapping pathways rather than serving compensatory roles. Notably, STX3 appears to have non-redundant functions, particularly in RPE cells, which require further experimental validation. Furthermore, our proteomic analysis identified a surprising array of novel STX1A and STX3 interactors with predominantly neuronal characteristics, including proteins implicated in neuronal survival, migration, development, and plasticity. These trafficking proteins exhibit both shared and unique binding partners, suggesting they carry out overlapping cellular functions while also maintaining distinct roles. Given the well-established epigenetic differences between RPE and neuronal cells, with RPE exhibiting a repressive chromatin state for photoreceptor-specific genes and a permissive state for neuronal-specific genes [57], the presence of these neuronal-associated proteins in RPE is particularly intriguing.

We found novel interactions of STX3 in ARPE19, including KIDINS220 [58-62], DNM1 [43,64], CHMP4B [65-66], CORO2B [67-69], THG1L [70-72], NME1 [73,74], and GNAO1 [75,76], which display strong neuronal specificity and have been linked to multiple neurological disorders. Thinning of the retinal layers contributed by the compromised RPE integrity also reflects the progression of neurodegenerative disorders [119]. Among the newly identified protein partners, including CORO2B, KIDINS220, and CHMP4B, also play a role in regulating actin dynamics and cellular plasticity. CORO2B, a focal adhesion protein, exerts a significant influence on actin dynamics via the cofilin-Rac1 pathway, thereby influencing cell mobility and migration [77]. In excitatory neurons, CORO2B regulates the excitatory-inhibitory balance, which is crucial for maintaining synaptic plasticity through a Rac1-dependent pathway. This, in turn, modulates actin dynamics, critical for synaptic strength regulation and neuronal migration [67,69]. Similarly, KIDINS220 plays a key role in initiating Rac1-mediated actin remodelling at neural tips, which is crucial for neurite outgrowth [59] and the establishment of neuronal polarity, which is achieved through the regulation of MAPK signalling [60,78] and the NGF-mediated secretory pathway [79]. CHMP4B also plays a regulatory role in Rac1 and Wnt signalling [65]. Its depletion was observed in Alzheimer’s disease [66], and it is vital for maintaining the structural integrity of the ciliary membrane [80]. Additionally, the Rac1 pathway at the presynaptic terminal is involved in vesicular priming and further regulates neurotransmitter release [81]. Notably, this pathway is influenced by CORO2B, KIDINS220, and CHMP4B, identified as interacting partners of STX3. The Rac1 pathway has a major role in maintaining synaptic plasticity as well as in actin dynamics. Additionally, it directly interacts with ZO1, the TJ protein, and Par3/Par6/aPKC protein complex in the formation, maintenance, and establishment of cell polarity [82]. Furthermore, KIDINS220-mediated PKA and NGF-dependent pathways activate calcium-CREB signalling, thereby regulating gene transcription involved in plasticity [79,83-88].

However, partners of STX1A have diverse functional specificities compared to STX3. PKN3 is a serine-threonine kinase, a cell adhesion protein involved in cytoskeletal organisation [89], specifically in adherens junctions [90], and negatively regulates RhoA signalling [91]. Notably, STX3-interacting proteins may directly modulate plasticity-related processes, while STX1A partner, PKN3, possibly enhances synaptic plasticity by negatively regulating RhoA signalling, thereby exerting a facilitatory effect on plastic changes. This suggests that STX3 and STX1A may converge on similar functional outcomes through the regulation of distinct, and potentially antagonistic pathways via their specific interacting partners, an interplay in a broader regulatory network which could be important for maintaining retinal homeostasis. Furthermore, SLC1A3, which is predominantly expressed in glia, is responsible for glutamate clearance, thereby reducing excitotoxicity due to excess glutamate [44, 92] is also a novel interactant of STX1A. Because glutamate plays a key part in visual signal processing, it stimulates light-evoked responses in the retina and is released by photoreceptors [115]. Since the role of SLC1A3 with STX1A is not yet reported in RPE, so, we speculate that STX1A-SLC1A3 interaction might also be functionally significant at the RPE-photoreceptor interface, facilitating clearance of excess glutamate from the photoreceptor by transporting it to RPE and thus maintaining retinal metabolic balance. Both STX1A and STX3 have several common interacting proteins specifically involved in protein movement and Golgi-mediated transport in RPE cells, such as KDELR2 [93] and NSF [94]. This overlap may suggest that while both the STXs contribute to vesicular trafficking processes in the RPE, each maintains a unique functional role that may be critical for preserving normal RPE and retinal physiology.

Interestingly, one of our observations from the initial molecular docking studies has revealed that the distinct and common interactants of STX1A and STX3 have different regions of interaction on the proteins. Specifically, these interactions occur at residues within the intrinsically disordered regions (IDRs) of the N-terminal domain (NTD), which overlap with the Sec1/Munc protein binding site (∼29-144), upstream of the H_abc_ domain (from∼1-28), in the H3 domain (∼189-259), and in the juxtamembrane region (∼258-282) (Sup. Table 4). Typically, the IDRs would provide conformational plasticity with relatively accessible multiple binding motifs where numerous partner proteins can bind with high specificity and average affinity, thereby altering their functional properties. These regions also serve as sites for post-translational modifications (PTMs) such as phosphorylation, which plays a crucial role in structural change and regulating signalling pathways [103-105,113,114]. In addition to this, numerous studies in rodents have documented that phosphorylation in the STX1A NTD, at Ser14 by CK2α [104] and Ser188 by DAP kinase [105], governs its binding dynamics with Mun18-1 and regulates neurotransmitter release and SNARE-mediated synaptic plasticity. Similarly, Bademosi et al. [106] demonstrated in *Drosophila* that STX1A is organised as a nanocluster and dynamically regulates the neurotransmitter release. A lysine-rich region within the H3 domain regulates this nanocluster formation, a process that is critical for presynaptic membrane organisation of SNAREs, specifically assembly/disassembly of SNARE complex [106]. Furthermore, previous reports have substantiated that the interaction of Munc18-1 to H_abc_ (∼29-144) and H3 (∼189-259) facilitates the conformational changes from open to closed structure so that the H_abc_ domain comes in proximity with the SNARE motif, and this inhibits SNARE complex assembly and subsequently blocks membrane fusion and synaptic exocytosis [101,102,107,108]. Notably, STX1A also interacts with Munc18-1 in the open state through the upstream region of H_abc_ in the NTD (∼1-28), while the H3 domain is dissociated from H_abc,_ which enables membrane fusion [108]. Remarkably, the H3 domain of STXs also displays dynamic structural flexibility, shifting between ordered and disordered conformations and functions as a regulatory switch that affects the participation of STXs in SNARE complex formation at the synapse. Further, the targeted mutation studies have confirmed that the mutation of hydrophobic residues in this region may impair the binding of the SNAP25 protein to the H3 domain, potentially disrupting synaptic transmission [107-112]. Taken together, the facts imply that the dynamic flexibility of these domains enables the precise assembly of the SNARE complex and optimises the efficiency of neurotransmitter release and SNARE-mediated synaptic plasticity, thereby adding a regulatory layer beyond basic on/off control mechanisms. To our surprise, our preliminary bioinformatics analysis revealed that the predicted interfacial amino acids in the interaction between novel interacting proteins and STXs reside within these described domains. We anticipate that the predicted unconventional binding of non-SNARE novel interactants identified through our LC-MS/MS analysis might be due to the dynamic nature of these domains, particularly in the NTD and H3 region, where binding was predominantly observed. This interaction could impart non-classical roles for STXs, such as SNARE-mediated synaptic plasticity, and could explain the binding of non-canonical proteins in the SNARE-interacting domains. However, these observations require thorough structural and experimental validation.

Retinal and RPE plasticity has been observed in amphibians, mice, and chicks, including their capacity for cell transdifferentiation, which facilitates repair and regeneration. However, this regenerative ability diminishes with cellular maturation in mice and chicks [50,95]. This report suggests that RPE cells possess an intrinsic self-repair capability under specific physiological conditions. Interestingly, in humans, RPE plasticity has been significantly observed in disease conditions such as proliferative vitreoretinopathy, where the mesenchymal transition occurs [96,97]. Amemiya et al. suggested that the presence of neuronal markers, such as Tuj1, in RPE cells might advocate their ability to adopt alternative cell fates [97]. This pleiotropic effect induced by Wnt7 also causes temporary changes in both the morphological and molecular structure of RPE [98,99], indicating the potential for plastic behaviour even in normal conditions. Therefore, we anticipate that the novel interactions of STX3 and STX1A in RPE cells might contribute to a static plasticity behaviour, which has not been observed in other epithelial cell types. This plasticity is essential for processes like the internalisation and phagocytosis of photoreceptor outer segments, as well as for self-repair and restoration. Hence, we propose that the identified novel interactions of STX3 and STX1A in RPE could act as an upstream regulatory event, driving actin remodelling and facilitating vesicular priming required for plasticity-related cellular processes. Given the neuroepithelial origin of RPE cells and their documented plasticity, particularly during development and under certain pathological conditions, our findings could suggest a broader role for RPE beyond its traditional photoreceptor support function [50, 98-100], which needs additional empirical validation.

Our study collectively confirms that STX3 is expressed in the human RPE and that its downregulation results in the destabilisation of TJs, indicating its potential role in preserving RPE cell polarity and structural integrity. Moreover, our data provides an initial indication that STX3 and STX1A interact with both distinct and common protein partners in the RPE, highlighting their functional significance. However, their specific interactions and roles within this context still need to be fully elucidated. The identification of proteins associated with neuronal plasticity within the STX3 and STX1A interactome may indicate their potential function in controlling RPE plasticity, thereby further emphasising their biological importance in maintaining retinal homeostasis. Nonetheless, the specific contribution of these interactions to RPE function and retinal physiology remains unclear and warrants in-depth investigation.

## Conflicts of Interest

The authors declare no conflict of interest.

## Data availability

All the data supporting the findings of the study are available within the article and supplementary information

## Authors contribution

The study was conceptualized, designed, and supervised by S. C. The research funding was acquired and administered by S. C. Experiments and the interpretation of the data were carried out by S. S; S. C; N. V. B and A. R. The manuscript was written by S. C., and A. R., and was reviewed with critical inputs by S. S. All authors read and approved the final manuscript.

## Acknowledgement

AR thanks the Indian Council of Medical Research (ICMR) for the SRF fellowship. CS the financial assistance provided by DBT-RGYI BT/PR15023/GBD/27/282/2010, Max Planck-India mobility grant ‘IGSTC/MPG/FS(SC)2012’, and the support from the University Grants Commission, New Delhi, for the award of Assistant Professorship under its Faculty Recharge Program (UGC-FRP). We would also like to acknowledge Prof. Reinhard Jahn, MPIBPC, Göttingen, for generously providing the STX1A and STX3 antibodies.

## Ethics approval

Donor eyes were procured from CU Shah Eye Bank, Sankara Nethralaya, Chennai, India, in accordance with the Institute Review Board, Vision Research Foundation, Chennai, strictly following the tenets of the Declaration of Helsinki. Informed consent was obtained for the use of donor eyes in accordance with ethical guidelines and institutional regulations.

## Declaration of generative AI and AI-assisted technologies in the writing process

During the preparation of the manuscript, we used AI-tools such as Trinka AI and Perplexity AI to improve the language and readability. After using these tools, we reviewed and edited the content as needed and took full responsibility for the publication’s content.

## Notes

### Competing Interest Statement

The authors have declared no competing interest.

## References

1. Bertolotti, E., Neri, A., Camparini, M., Macaluso, C., & Marigo, V. (2014). Stem cells as source for retinal pigment epithelium transplantation. In Progress in Retinal and Eye Research (Vol. 42).

2. Strauss O. The Retinal Pigment Epithelium in Visual Function. Physiological Reviews. 2005.

3. Lehmann GL, et al., Plasma membrane protein polarity and trafficking in RPE cells: past, present and future. Exp Eye Res. 2014.

4. Finnemann SC, Bonilha VL, Marmorstein AD, Rodriguez-Boulan E. Phagocytosis of rod outer segments by retinal pigment epithelial cells requires alpha(v)beta5 integrin for binding but not for internalization. Proc Natl Acad Sci U S A. 1997;94(24):12932–12937.

5. Caceres, P. S., & Rodriguez-Boulan, E. (2020). Retinal pigment epithelium polarity in health and blinding diseases. In Current Opinion in Cell Biology (Vol. 62, pp. 37–45). Elsevier Ltd.

6. Marmorstein, A. D. (2001). The polarity of the retinal pigment epithelium. In Traffic (Vol. 2, Issue 12, pp. 867–872).

7. Pinilla, I., Keeley, P. W., & Lillo, C. (n.d.). Scribble basal polarity acquisition in RPE cells and its mislocalization in a pathological AMD-like model.

8. Hoo, L. S., Banna, C. D., Radeke, C. M., Sharma, N., Albertolle, M. E., Low, S. H., Weimbs, T., & Vandenberg, C. A. (2016). The SNARE protein syntaxin 3 confers specificity for polarized axonal trafficking in neurons. PLoS ONE, 11(9).

9. Li, X., Low, H., Miura, M., & Weimbs, T. (2002). SNARE expression and localization in renal epithelial cells suggest mechanism for variability of trafficking phenotypes.

10. Low, S. H., Marmorstein, L. Y., Miura, M., Li, X., Kudo, N., Marmorstein, A. D., & Weimbs, T. (2002). Retinal pigment epithelial cells exhibit unique expression and localization of plasma membrane syntaxins which may contribute to their trafficking phenotype. In Journal of Cell Science (Vol. 115, Issue 23, pp. 4545–4553).

11. Low, S. H., Vasanji, A., Nanduri, J., He, M., Sharma, N., Koo, M., Drazba, J., & Weimbs, T. (2006). Syntaxins 3 and 4 are concentrated in separate clusters on the plasma membrane before the establishment of cell polarity. Molecular Biology of the Cell, 17(2), 977–989.

12. Sharma, N., Low, S. H., Misra, S., Pallavi, B., & Weimbs, T. (2006). Apical targeting of syntaxin 3 is essential for epithelial cell polarity. Journal of Cell Biology, 173(6), 937–948.

13. Zhou Q, Xiao J, Liu Y. Participation of syntaxin 1A in membrane trafficking involving neurite elongation and membrane expansion. J Neurosci Res. 2000;61(3):321–328.

14. Bin, N. R., Huang, M., & Sugita, S. (2019). Investigating the Role of SNARE Proteins in Trafficking of Postsynaptic Receptors using Conditional Knockouts. In Neuroscience (Vol. 420).

15. Sutton, R. B., Fasshauer, D., Jahn, R., & Brunger, A. T. (1998). Crystal structure of a SNARE complex involved in synaptic exocytosis at 2.4 Å resolution. In Nature (Vol. 395, Issue 6700).

16. Lafont F, Verkade P, Galli T, Wimmer C, Louvard D, Simons K. Raft association of SNAP receptors acting in apical trafficking in Madin-Darby canine kidney cells. Proc Natl Acad Sci U S A. 1999;96(7):3734–3738.

17. Low SH, Chapin SJ, Wimmer C, et al. The SNARE machinery is involved in apical plasma membrane trafficking in MDCK cells. J Cell Biol. 1998;141(7):1503–1513.

18. Hayashi K, et al., Role of syntaxin3 an apical polarity protein in poorly polarized keratinocytes: regulation of asymmetric barrier formations in the skin epidermis. Cell Tissue Res. 2023.

19. Quiñones B, Riento K, Olkkonen VM, Hardy S, Bennett MK. Syntaxin 2 splice variants exhibit differential expression patterns, biochemical properties and subcellular localizations. J Cell Sci. 1999;112 (Pt 23):4291

20. Kakakhel M, et al., Syntaxin 3 is essential for photoreceptor outer segment protein trafficking and survival. Proc Natl Acad Sci. 2020.

21. Perez-Hurtado M, et al., Syntaxin 3 is haplosufficient for long-term photoreceptor survival in the mouse retina. Front Ophthalmol (Lausanne). 2023.

22. Gething, C., Ferrar, J., Misra, B., Howells, G., Andrzejewski, A. L., Bowen, M. E., & Choi, U. B. (2022). Conformational change of Syntaxin-3b in regulating SNARE complex assembly in the ribbon synapses. Scientific Reports, 12(1).

23. Weber T, Zemelman BV, McNew JA, et al. SNAREpins: minimal machinery for membrane fusion. Cell. 1998;92(6):759–772.

24. Curtis LB, Doneske B, Liu X, Thaller C, McNew JA, Janz R. Syntaxin 3b is a t-SNARE specific for ribbon synapses of the retina. J Comp Neurol. 2008;510(5):550–559.

25. Curtis L, Datta P, Liu X, Bogdanova N, Heidelberger R, Janz R. Syntaxin 3B is essential for the exocytosis of synaptic vesicles in ribbon synapses of the retina. Neuroscience. 2010;166(3):832–841.

26. Nishad R, Betancourt-Solis M, Dey H, Heidelberger R, McNew JA. Regulation of Syntaxin3B-Mediated Membrane Fusion by T14, Munc18, and Complexin. Biomolecules. 2023;13(10):1463. Published 2023 Sep 28.

27. Zhu, M., Provis, J. M., & Penfold, P. L. (1998). Isolation, culture and characteristics of human foetal and adult retinal pigment epithelium. Australian and New Zealand Journal of Ophthalmology, 26(SUPPL. 1).

28. Palanisamy, K., Karunakaran, C., Raman, R., & Chidambaram, S. (2019). Optimization of an in vitro bilayer model for studying the functional interplay between human primary retinal pigment epithelial and choroidal endothelial cells isolated from donor eyes. BMC Research Notes, 12(1).

29. Day, P., Riggs, K. A., Hasan, N., Corbin, D., Humphrey, D., & Hu, C. (2011). Syntaxins 3 and 4 mediate vesicular trafficking of α5β1 and α3β1 integrins and cancer cell migration. International Journal of Oncology, 39(4).

30. Pathan, M., Keerthikumar, S., Ang, C. S., Gangoda, L., Quek, C. Y. J., Williamson, N. A., Mouradov, D., Sieber, O. M., Simpson, R. J., Salim, A., Bacic, A., Hill, A. F., Stroud, D. A., Ryan, M. T., Agbinya, J. I., Mariadason, J. M., Burgess, A. W., & Mathivanan, S. (2015). FunRich: An open access standalone functional enrichment and interaction network analysis tool. Proteomics, 15(15).

31. Webb B, Sali A. Comparative Protein Structure Modeling Using MODELLER. Curr Protoc Bioinformatics. 2016.

32. Webb B, Sali A. Protein Structure Modeling with MODELLER. Methods Mol Biol. 2021.

33. Roche DB, et al., The IntFOLD server: an integrated web resource for protein fold recognition, 3D model quality assessment, intrinsic disorder prediction, domain prediction and ligand binding site prediction. Nucleic Acids Res. 2011.

34. Lovell SC, et al., Structure validation by Calpha geometry: phi,psi and Cbeta deviation. Proteins. 2003.

35. Wiederstein M, Sippl MJ. ProSA-web: interactive web service for the recognition of errors in three-dimensional structures of proteins. Nucleic Acids Research. 2007.

36. Sippl MJ. Recognition of errors in three-dimensional structures of proteins. Proteins. 1993.

37. Linding R, et al., Protein disorder prediction: implications for structural proteomics. Structure. 2003.

38. Honorato RV, et al. The HADDOCK2.4 web server for integrative modeling of biomolecular complexes. Nat Protoc. 2024.

39. Venkatraman V, Yang YD, Sael L, Kihara D. Protein-protein docking using region-based 3D Zernike descriptors. BMC Bioinformatics. 2009;10:407. Published 2009 Dec 9.

40. Laskowski RA, et al., PDBsum: Structural summaries of PDB entries. Protein Science: A Publication of the Protein Society. 2017.

41. E. Krissinel and K. Henrick, PDBePISA: Inference of macromolecular assemblies from crystalline state, J. Mol. Biol. 2007.

42. Tebbe L, et al., The role of syntaxins in retinal function and health. Front Cell Neurosci. 2024.

43. Noakes PG, et al., Expression and localisation of dynamin and syntaxin during neural development and neuromuscular synapse formation. J Comp Neurol. 1999

44. Yu Y-X, et al., Syntaxin 1A promotes the endocytic sorting of EAAC1 leading to inhibition of glutamate transport. Journal of Cell Science. 2006

45. Dos Santos B, et al., The impact of purine nucleosides on neuroplasticity in the adult brain. Purinergic Signal. 2024.

46. Yan J, et al., Disrupted de novo pyrimidine biosynthesis impairs adult hippocampal neurogenesis and cognition in pyridoxine-dependent epilepsy. Sci Adv. 2024.

47. Moro A, van Nifterick A, Toonen RF, Verhage M. Dynamin controls neuropeptide secretion by organizing dense-core vesicle fusion sites. Sci Adv. 2021;7(21):eabf0659. doi:10.1126/sciadv.abf0659

48. Galas M-C, et al., Presence of Dynamin—Syntaxin Complexes Associated with Secretory Granules in Adrenal Chromaffin Cells. J Neurochem. 2000.

49. Bennis, A., Gorgels, T. G. M. F., Ten Brink, J. B., Van Der Spek, P. J., Bossers, K., Heine, V. M., & Bergen, A. (2015). Comparison of mouse and human retinal pigment epithelium gene expression profiles: Potential implications for age-related macular degeneration. PLoS ONE, 10(10).

50. Fuhrmann S, Zou C, Levine EM. Retinal pigment epithelium development, plasticity, and tissue homeostasis, Exp Eye Res. 2014

51. Gupta, S., Lytvynchuk, L., Ardan, T., Studenovska, H., Faura, G., Eide, L., Znaor, L., Erceg, S., Stieger, K., Motlik, J., Bharti, K., & Petrovski, G. (2023). Retinal Pigment Epithelium Cell Development: Extrapolating Basic Biology to Stem Cell Research. In Biomedicines (Vol. 11, Issue 2).

52. Swaroop, A., Kim, D., & Forrest, D. (2010). Transcriptional regulation of photoreceptor development and homeostasis in the mammalian retina. In Nature Reviews Neuroscience (Vol. 11, Issue 8).

53. Pournami F, Mk AK, Panackal AV, Nandakumar A, Prabhakar J, Jain N. Microvillus Inclusion Disease: A Rare Mutation of STX3 in Exon 9 Causing Fatal Congenital Diarrheal Disease. J Pediatr Genet. 2020;11(2):154–157..

54. Vardar G, Chang S, Arancillo M, Wu YJ, Trimbuch T, Rosenmund C. Distinct Functions of Syntaxin-1 in Neuronal Maintenance, Synaptic Vesicle Docking, and Fusion in Mouse Neurons. J Neurosci. 2016;36(30):7911–7924.

55. Shi S, Ma K, Bin NR, et al. Syntaxin-3 is dispensable for basal neurotransmission and synaptic plasticity in postsynaptic hippocampal CA1 neurons. Sci Rep. 2020;10(1):709.

56. Vardar G, Salazar-Lázaro A, Brockmann M, et al. Reexamination of N-terminal domains of syntaxin-1 in vesicle fusion from central murine synapses. Elife. 2021;10:e69498.

57. Dvoriantchikova G, Seemungal RJ, Ivanov D. DNA Methylation Dynamics During the Differentiation of Retinal Progenitor Cells Into Retinal Neurons Reveal a Role for the DNA Demethylation Pathway. Front Mol Neurosci. 2019;12:182.

58. Arévalo JC, Wu SH, Takahashi T, et al. The ARMS/Kidins220 scaffold protein modulates synaptic transmission. Mol Cell Neurosci. 2010;45(2):92–100.

59. Neubrand VE, et al., Kidins220/ARMS regulates Rac1-dependent neurite outgrowth by direct interaction with the RhoGEF Trio. J Cell Sci. 2010.

60. Higuero AM, et al., Kidins220/ARMS modulates the activity of microtubule-regulating proteins and controls neuronal polarity and development. J Biol Chem. 2010.,

61. Zhang J, et al., Gain-of-Function KIDINS220 Variants Disrupt Neuronal Development and Cause Cerebral Palsy. Mov Disord. 2024.,

62. Scholz-Starke J, et al., Kidins220/ARMS Is a Novel Modulator of Short-Term Synaptic Plasticity in Hippocampal GABAergic Neurons, PLoS One. 2012

63. Galas MC, Chasserot-Golaz S, Dirrig-Grosch S, Bader MF. Presence of dynamin--syntaxin complexes associated with secretory granules in adrenal chromaffin cells. J Neurochem. 2000 Oct;75(4):1511–9.

64. Nakashima M, et al., De novo DNM1 mutations in two cases of epileptic encephalopathy. Epilepsia. 2016.

65. Fumoto K, et al., Wnt5a signaling controls cytokinesis by correctly positioning ESCRT-III at the midbody. J Cell Sci. 2012.

66. Ding Y, et al., Pyroptosis Signature Gene CHMP4B Regulates Microglia Pyroptosis by Inhibiting GSDMD in Alzheimer’s Disease. Mol Neurobiol. 2024.

67. Chen Y, et al., Coronin 2B Regulates Neuronal Migration via Rac1-Dependent Multipolar-Bipolar Transition. J Neurosci. 2023.

68. Wu H, et al., Coronin 2B deficiency induces nucleolar stress and neuronal apoptosis. Cell Death Dis. 2024.

69. Chen Y, et al., Coronin 2B regulates dendrite outgrowth by modulating actin dynamics FEBS Lett. 2020.

70. Han R, et al., Compound heterozygous variants of THG1L result in autosomal recessive cerebellar ataxia. J Hum Genet. 2023.

71. Edvardson S, et al., A mutation in the THG1L gene in a family with cerebellar ataxia and developmental delay. Neurogenetics. 2016.

72. Rabin R, et al., Severe epileptic encephalopathy associated with compound heterozygosity of THG1L variants in the Ashkenazi Jewish population. Am J Med Genet A. 2021.

73. Anantha J, et al., STRAP and NME1 Mediate the Neurite Growth-Promoting Effects of the Neurotrophic Factor GDF5. iScience. 2020.

74. Anantha J, et al., NME1 Protects Against Neurotoxin-, α-Synuclein- and LRRK2-Induced Neurite Degeneration in Cell Models of Parkinson’s Disease. Mol Neurobiol. 2022.

75. Akamine S, et al. GNAO1 organizes the cytoskeletal remodeling and firing of developing neurons. FASEB J. 2020.

76. Saitsu H, et al., Phenotypic spectrum of GNAO1 variants: epileptic encephalopathy to involuntary movements with severe developmental delay. Eur J Hum Genet. 2016.

77. Wu H, et al., Coronin 2B deficiency induces nucleolar stress and neuronal apoptosis. Cell Death Dis. 2024.

78. Bracale A, et al., Kidins220/ARMS Is Transported by a Kinesin-1–based Mechanism Likely to be Involved in Neuronal Differentiation Molecular Biology of the Cell. Mol Biol Cell. 2007.

79. López-Benito S, et al., ARMS/Kidins220 and synembryn-B levels regulate NGF-mediated secretion. J CellSci. 2016.

80. Jung E, et al., ESCRT subunit CHMP4B localizes to primary cilia and is required for the structural integrity of the ciliary membrane. FASEB J. 2020.

81. Keine C, et al., Presynaptic Rac1 controls synaptic strength through the regulation of synaptic vesicle priming. Elife. 2022

82. Gopalakrishnan S, Hallett MA, Atkinson SJ, Marrs JA. aPKC-PAR complex dysfunction and tight junction disassembly in renal epithelial cells during ATP depletion. Am J Physiol Cell Physiol. 2007;292(3):C1094–C1102.

83. Finsterwald C, et al., Regulation of dendritic development by BDNF requires activation of CRTC1 by glutamate. J Biol Chem. 2010

84. Tao X, et al., Ca2+ Influx Regulates BDNF Transcription by a CREB Family Transcription Factor-Dependent Mechanism. Neuron. 1998

85. López-Benito S, et al., Regulation of BDNF Release by ARMS/Kidins220 through Modulation of Synaptotagmin-IV Levels. J Neurosci. 2018

86. Waltereit R, Weller M. Signaling from cAMP/PKA to MAPK and synaptic plasticity. Mol Neurobiol. 2003

87. Alberini CM. Transcription Factors in Long-Term Memory and Synaptic Plasticity. Physiol Rev. 2009.

88. Riccio A, et al., Mediation by a CREB family transcription factor of NGF-dependent survival of sympathetic neurons. Science. 1999

89. Mukai H, Muramatsu A, Mashud R, et al. PKN3 is the major regulator of angiogenesis and tumor metastasis in mice. Sci Rep. 2016;6:18979.

90. Kristin Möpert, Kathrin Löffler, Nadine Röder, Jörg Kaufmann, Ansgar Santel, Depletion of protein kinase N3 (PKN3) impairs actin and adherens junctions dynamics and attenuates endothelial cell activation, European Journal of Cell Biology, Volume 91, Issue 9, 2012, Pages 694–705

91. Dibus M, Brábek J, Rösel D. A Screen for PKN3 Substrates Reveals an Activating Phosphorylation of ARHGAP18. Int J Mol Sci. 2020;21(20):7769.

92. Gorostiola González, M., Sijben, H. J., Dall’ Acqua, L., Liu, R., IJzerman, A. P., Heitman, L. H., & van Westen, G. J. P. (2023). Molecular insights into disease-associated glutamate transporter (EAAT1 / SLC1A3) variants using in silico and in vitro approaches. Frontiers in Molecular Biosciences, 10.

93. Wires ES, Trychta KA, Kennedy LM, Harvey BK. The Function of KDEL Receptors as UPR Genes in Disease. Int J Mol Sci. 2021;22(11):5436.

94. Yang J, Kong L, Zou L, Liu Y. The role of synaptic protein NSF in the development and progression of neurological diseases. Front Neurosci. 2024;18:1395294.

95. Sharma P, Ramachandran R. Retina regeneration: lessons from vertebrates. Oxf Open Neurosci. 2022;1:kvac012.

96. Salero E, et al., Adult Human RPE Can Be Activated into a Multipotent Stem Cell that Produces Mesenchymal Derivatives. Cell Stem Cell. 2012.

97. Amemiya K, et al., Adult human retinal pigment epithelial cells capable of differentiating into neurons. Biochem Biophys Res Commun. 2004.

98. Kuznetsova AV, et al., Plasticity of adult human retinal pigment epithelial cells. Int J Clin Exp Med 2016

99. Lueck K, et al., Annexin A8 regulates Wnt signaling to maintain the phenotypic plasticity of retinal pigment epithelial cells. Sci Rep. 2020.

100. Milyushina LA, et al., Phenotypic Plasticity of Retinal Pigment Epithelial Cells from Adult Human Eye In Vitro. Bull Exp Biol Med. 2011.

101. Ma C, Li W, Xu Y, Rizo J. Munc13 mediates the transition from the closed syntaxin-Munc18 complex to the SNARE complex. Nat Struct Mol Biol. 2011;18(5):542–549.

102. Bin NR, Jung CH, Kim B, et al. Chaperoning of closed syntaxin-3 through Lys46 and Glu59 in domain 1 of Munc18 proteins is indispensable for mast cell exocytosis. J Cell Sci. 2015;128(10):1946–1960.

103. Chakrabarti P, Chakravarty D. Intrinsically disordered proteins/regions and insight into their biomolecular interactions. Biophys Chem. 2022;283:106769.

104. Shi VH, Craig TJ, Bishop P, et al. Phosphorylation of Syntaxin-1a by casein kinase 2α regulates pre-synaptic vesicle exocytosis from the reserve pool. J Neurochem. 2021;156(5):614–623.

105. Tian JH, Das S, Sheng ZH. Ca2+-dependent phosphorylation of syntaxin-1A by the death-associated protein (DAP) kinase regulates its interaction with Munc18. J Biol Chem. 2003;278(28):26265–26274.

106. Bademosi, A., Lauwers, E., Padmanabhan, P. et al. In vivo single-molecule imaging of syntaxin1A reveals polyphosphoinositide- and activity-dependent trapping in presynaptic nanoclusters. Nat Commun 7, 13660 (2016).

107. Zilly FE, Sørensen JB, Jahn R, Lang T. Munc18-bound syntaxin readily forms SNARE complexes with synaptobrevin in native plasma membranes. PLoS Biol. 2006;4(10):e330.

108. Dawidowski D, Cafiso DS. Allosteric control of syntaxin 1a by Munc18-1: characterization of the open and closed conformations of syntaxin. Biophys J. 2013;104(7):1585–1594.

109. Fergestad T, Wu MN, Schulze KL, Lloyd TE, Bellen HJ, Broadie K. Targeted mutations in the syntaxin H3 domain specifically disrupt SNARE complex function in synaptic transmission. J Neurosci. 2001;21(23):9142–9150.

110. Yang X, Tu W, Gao X, Zhang Q, Guan J, Zhang J. Functional regulation of syntaxin-1: An underlying mechanism mediating exocytosis in neuroendocrine cells. Front Endocrinol (Lausanne). 2023;14:1096365. Published 2023 Jan 19.

111. Li G, Yang Q, Alexander EA, Schwartz JH. Syntaxin 1A has a specific binding site in the H3 domain that is critical for targeting of H+-ATPase to apical membrane of renal epithelial cells. Am J Physiol Cell Physiol. 2005;289(3):C665–C672.

112. Zhong P, Chen YA, Tam D, Chung D, Scheller RH, Miljanich GP. An alpha-helical minimal binding domain within the H3 domain of syntaxin is required for SNAP-25 binding. Biochemistry. 1997;36(14):4317–4326. doi:10.1021/bi9625408

113. Wright, P., Dyson, H. Intrinsically disordered proteins in cellular signalling and regulation. Nat Rev Mol Cell Biol 16, 18–29 (2015).

114. Moses, D., Guadalupe, K., Yu, F. et al. Structural biases in disordered proteins are prevalent in the cell. Nat Struct Mol Biol 31, 283–292 (2024).

115. Niklaus S, Glasauer SMK, Kovermann P, et al. Glutamate transporters are involved in direct inhibitory synaptic transmission in the vertebrate retina. Open Biol. 2024;14(7):240140. doi:10.1098/rsob.240140

116. Shi L, Kümmel D, Coleman J, Melia TJ, Giraudo CG. Dual roles of Munc18-1 rely on distinct binding modes of the central cavity with Stx1A and SNARE complex. Mol Biol Cell. 2011 Nov;22(21):4150–60. doi: 10.1091/mbc.E11-02-0150. Epub 2011 Sep 7. PMID: 21900493; PMCID: PMC3204075.

117. Hackmann Y, Graham SC, Ehl S, Höning S, Lehmberg K, Aricò M, Owen DJ, Griffiths GM. Syntaxin binding mechanism and disease-causing mutations in Munc18-2. Proc Natl Acad Sci U S A. 2013 Nov 19;110(47):E4482–91. doi: 10.1073/pnas.1313474110.

118. Brochetta C, Suzuki R, Vita F, et al. Munc18-2 and syntaxin 3 control distinct essential steps in mast cell degranulation. J Immunol. 2014;192(1):41–51. doi:10.4049/jimmunol.1301277.

119. Uchida A, Pillai JA, Bermel R, Jones SE, Fernandez H, Leverenz JB, Srivastava SK, Ehlers JP. Correlation between brain volume and retinal photoreceptor outer segment volume in normal aging and neurodegenerative diseases. PLoS One. 2020 Sep 3;15(9):e0237078. doi: 10.1371/journal.pone.0237078.

