## Supplementary data for "Syntaxin-3 regulates Tight Junction Assembly in Human Retinal Pigment Epithelium"

**Table 1. List of primer sequences used in q-RT-PCR**

| No | Gene | Forward primer | Reverse Primer |
|---|---|---|---|
| 1 | GAPDH | gaaggtcggagtcaacggatt | cgctcctggaagatggtga |
| 2 | 18S rRNA | aacccgttgaaccccatt | ccatccaatcggtagtagcg |
| 3 | STX1A | gaccgcttcatggatgagtt | cttggaacgaactttgtttgc |
| 4 | STX2 | ggaagaatgatgatggagac | agaaagaatgatgctgtgg |
| 5 | STX3 variant1 | attggactttccgttgggct | acattcccccagggctagat |
| 6 | STX3 variant2 | aggcccggaagaaactgattt | acattcccccagggctagat |
| 7 | STX4 | tgagctgcacgacatattca | tgtgcgacattatccaacca |
| 8 | STX6 | ggtacagaaagcagtcaacac | tcatcaaggtcctctagatcc |
| 9 | STX7 | aacacctcaagattcacctg | gttgtgaactctgccactaag |
| 10 | STX8 | aagcttaccgtgacaatcag | aaggatgccagaagtagtctc |
| 11 | STX12 | atgagccagctaggaactaag | gttccttctgaagtctctgct |
| 12 | STX16 | cagcttagatccagaagcag | ctgagtgatctcttgggtagtt |
| 13 | SNAP23 | ggacatgagagagacagagaa | gcctggctgtttagatactac |
| 14 | SNAP25 | atgctggtatcaggactttg | tgttacagggacacacacaa |
| 15 | SNAP29 | agaacaggaagcaaagtacc | ctgtcgatcttctggtgatag |
| 16 | VAMP1 | acagacgactacagcaaacc- | ttttccaccaatacttcctc |
| 17 | VAMP2 | ctccaaacctcaccagtaac | ggtttttccaccagtatttg |
| 18 | VAMP3 | aatcgaagacttcagcagac | cagagagcttctggtctcttt |
| 19 | VAMP7 | cgagttctcaagtgtcttagc | agcaagatttctgctggtag |
| 20 | VAMP8 | gaggtggagggagttaagaat | acaatcacgcagataaggac |

| | | | |
|---|---|---|---|
| 21 | Rab1 | gttacttctgattggcgact | catgggctcctctgtaataa |
| 22 | Rab3 | cggatcagaacttcgactac | agcttgatcctcttgtcgt |
| 23 | Rab5 | atacagctggtcaagaacga | ttaggacttgcttgcctct |
| 24 | Rab6 | tagcctcattcccagttaca | cacttcctctttctgttctgac |
| 25 | Rab8 | atcacaacggcctactacag | ccttggaaacttgtctcttg |
| 26 | Rab11 | gagctgtaggtgccttattg | gctcttgcttcatctgtagg |

**Table 2. List of antibodies used for Western blot analysis and immunofluorescence assay**

| **No** | **Antibody** | **Company** |
|---|---|---|
| 1 | Syntaxin3 | Synaptic systems |
| 2 | Syntaxin1A | Synaptic systems |
| 3 | β-Actin | Santa Cruz Biotechnology |
| 4 | Occludin | Invitrogen |
| 5 | ZO1 | Invitrogen |

**Sup.Fig 1. Raw data for agarose gel images and the Western blot**

**a**

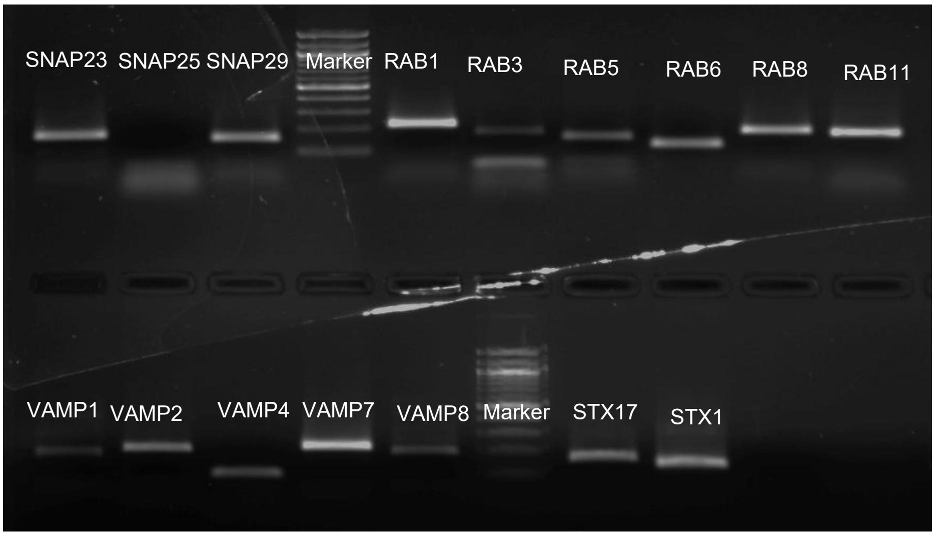


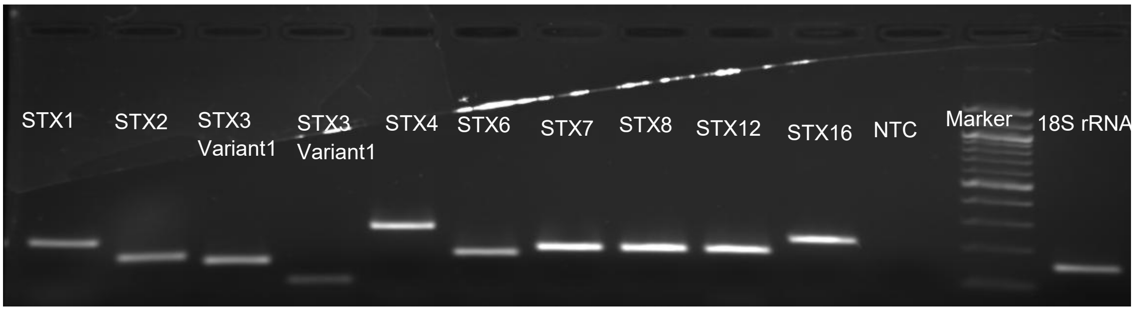


**b**

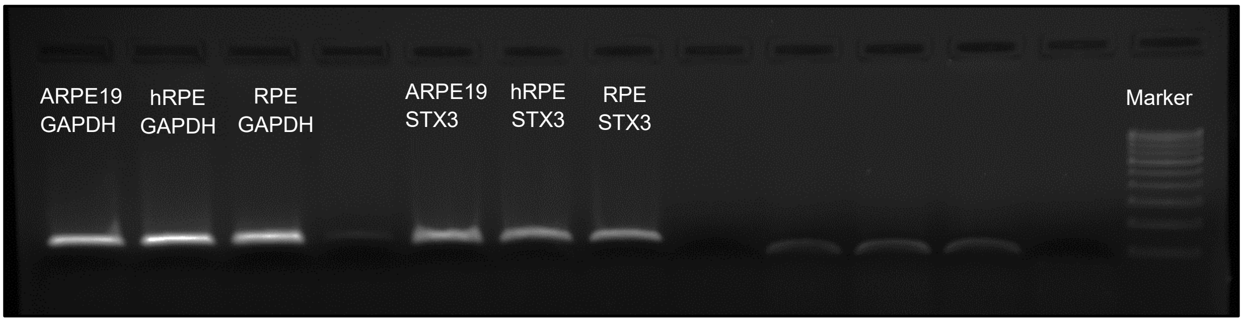

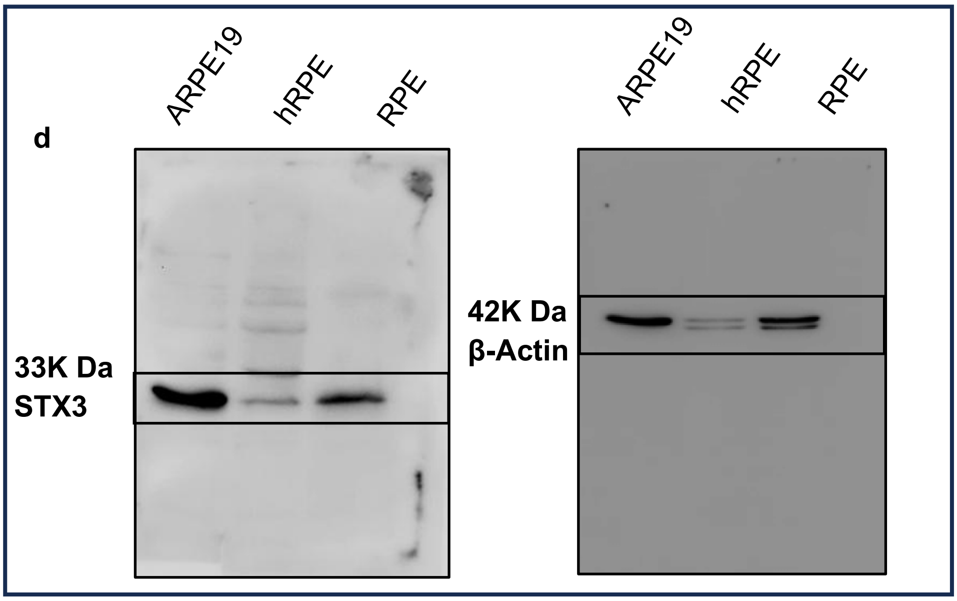


**Fig.1 Identification and validation of STX3 expression in RPE**: **a**: The RT-PCR results depict the expression of various SNARE transcripts, Syntaxins 1A, 2, 3 (variant1 and 2), 4, 6, 7, 8, 12 and 16; SNAP 23,25 and 29; VAMPs 1, 2, 3, 7 and 8; Rabs 1, 3, 5, 6, 8 and 11 in ARPE19 cell line. **b**: Variants of STX3 were detected in ARPE19, primary RPE cells, and donor RPE tissue. **d**: STX3 protein expression in ARPE19, primary RPE cells, and donor RPE tissue was validated by Western blot where β-Actin was used as a loading control.

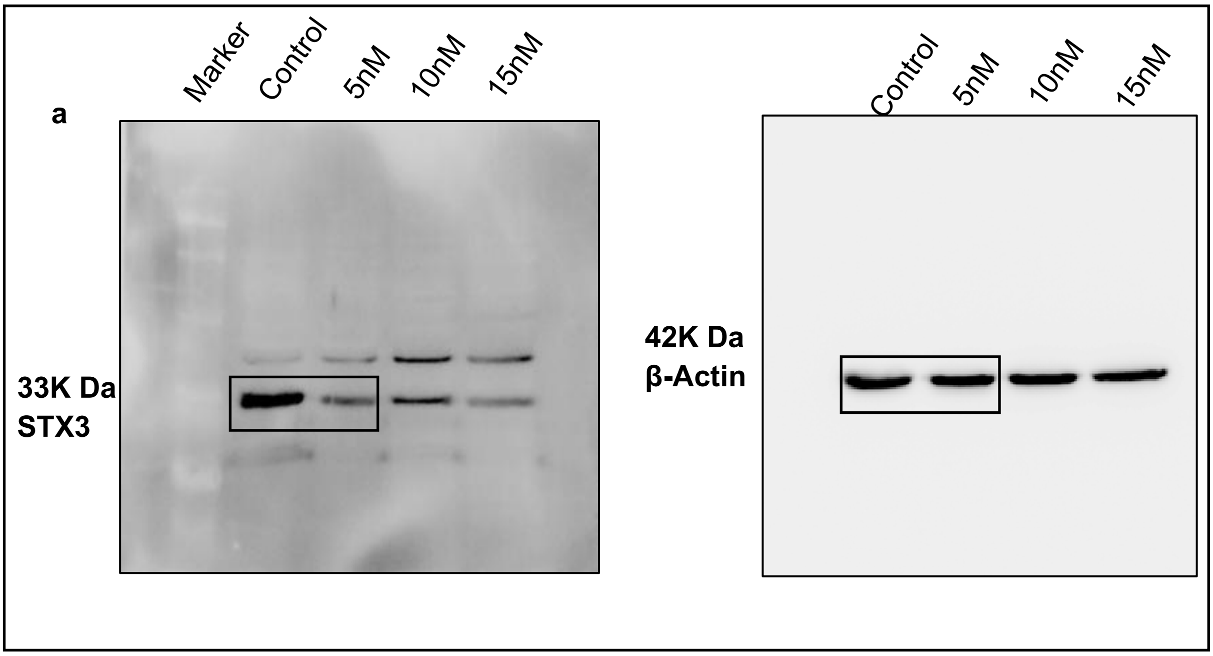


**Sup. Fig.2**: **The Western blot analysis showing siRNA-mediated knockdown of STX3 in ARPE19 cells:** si-STX3 was used at 5nM, 10nM and 15nM concentrations to knockdown STX3 in ARPE19. The blot was stripped and reprobed for the loading control β-Actin

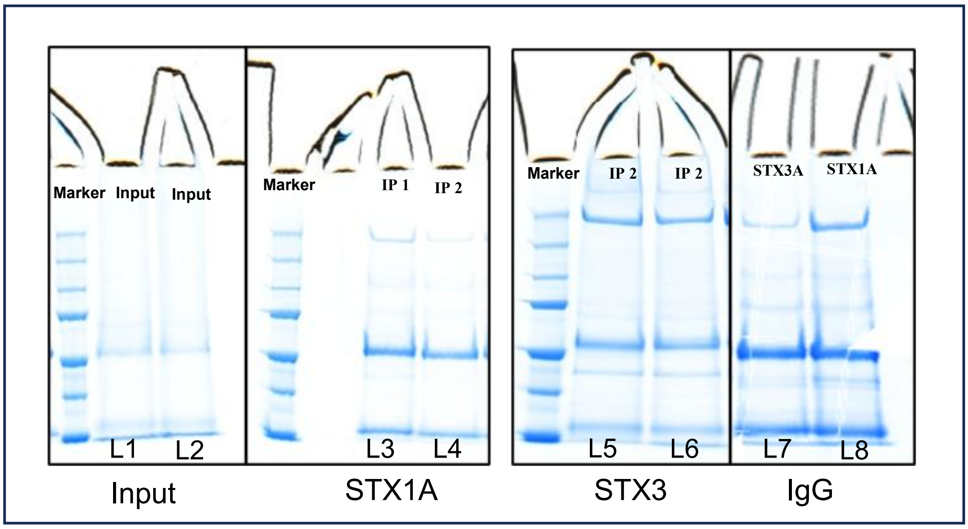


**Sup. Fig.3: Coomassie-stained SDS-PAGE gel of the immunoprecipitated samples.** ARPE19 cell lysate was subjected to immunoprecipitation using Anti-Rabbit STX3 and Anti-Rabbit STX1A antibodies, followed by SDS-PAGE analysis. Immunoprecipitation samples of STX1A (L3, L4) and STX3 (L5, L6) Input (L1,L2), IgG control (L7, L8).

**Table. 3a:** The Haddock results for docking studies involving STX3 with its novel and distinct interacting partners

| | **STX3 & DNM1** | **STX3 & CHMP4B** | **STX3 & CORO2B** | **STX3 & KIDINS220** | **STX1A & DNM1** |
|---|---|---|---|---|---|
| **Haddock Score** | -101.2±5.2 | -132±16.7 | -100±18.1 | -75.4±24.1 | -53.9+/-4.9 |
| **Buried surface area** | 3156.6±233.7 | 3411.9±206 | 4897.7±136.6 | 3441.4±202.8 | 2316.8+/-127.7 |
| **Z Score** | -1.3 | -2-2 | -2.2 | -1.7 | -2.2 |
| **Hydrogen bonds** | 15 | 16 | 18 | 10 | 9 |
| **Salt bridge** | 3 | 7 | 6 | 4 | 1 |
| **Electrostatic energy** | -379.4±54.3 | -591.6±71.5 | -420.3±83.7 | -496.6±53.4 | -261.6±28.4 |
| **Vanderwaals force** | -87.5±10.9 | -83.4±13.3 | -153.9±9.2 | -94.4±9.7 | -80.0+/-7.2 |

**Table. 3b:** The LZerD results for docking studies involving STX3 and STX1A with its novel and distinct interacting partners

| **No** | **STX-Interactant** | **GOAP Score** | **DFIRE Score** | **ITScore** | **Ranksum Score** |
|---|---|---|---|---|---|
| 1 | STX1A-SLC1A3 | -92218.69 | -71345.00 | -37081.21 | 424 |
| 2 | STX1A-NSF | -147748.88 | -85744.94 | -44174.68 | 2070 |
| 3 | STX1A-AMBP | -78984.40 | -50213.08 | -27113.41 | 62 |
| 4 | STX1A-KDELR2 | -69063.56 | -48770.75 | -26548.02 | 14 |
| 5 | STX1A-LRG1 | -89196.42 | -55918.11 | -28225.44 | 37 |
| 6 | STX1A-PKN3 | -144991.87 | -96939.56 | -50201.93 | 14 |
| 7 | STX3-AMBP | -78984.40 | 50213.08 | -27113.41 | 62 |
| 8 | STX3-NSF | -157507.02 | -91236.13 | -47229.30 | 14 |
| 9 | STX3-LRG1 | -89196.42 | -55918.11 | -28225.44 | 37 |
| 10 | STX3-KDELR2 | -69063.56 | -48770.75 | -26548.02 | 14 |

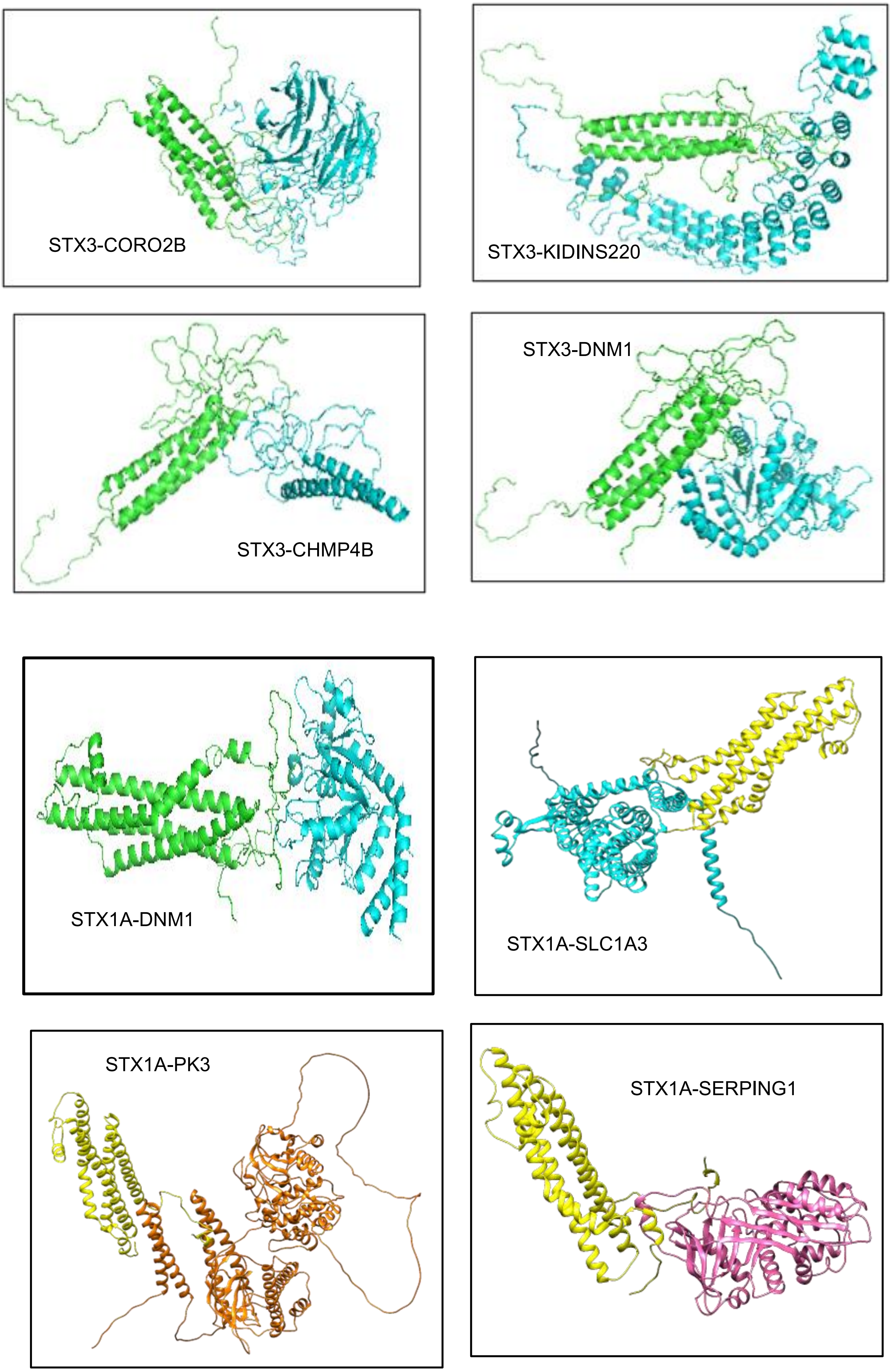
STX3-CORO2B
STX3-KIDINS220
STX3-CHMP4B
STX3-DNM1
STX1A-DNM1
STX1A-SLC1A3
STX1A-PK3
STX1A-SERPING1

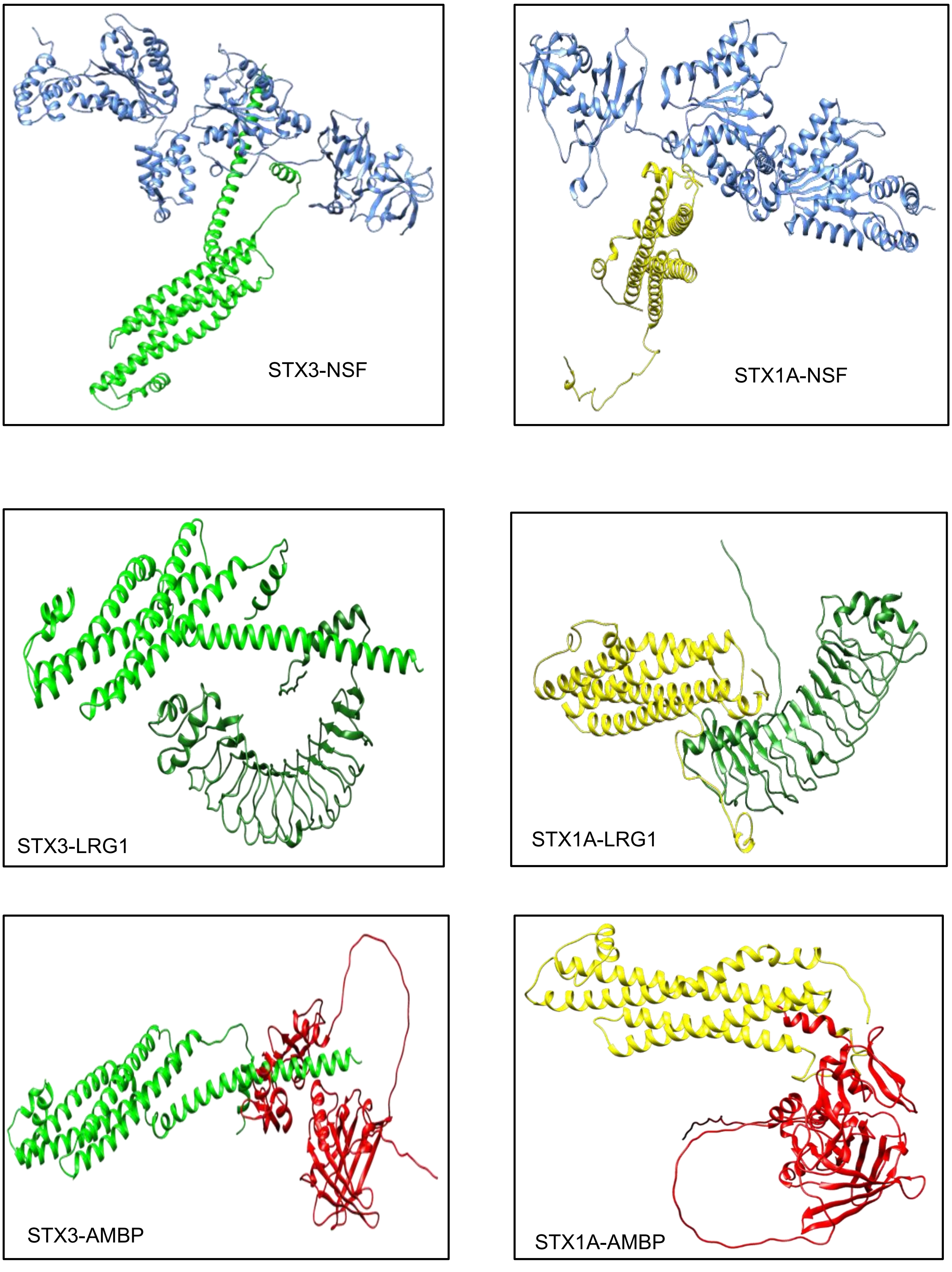
STX3-NSF
STX1A-NSF
STX3-LRG1
STX1A-LRG1
STX3-AMBP
STX1A-AMBP

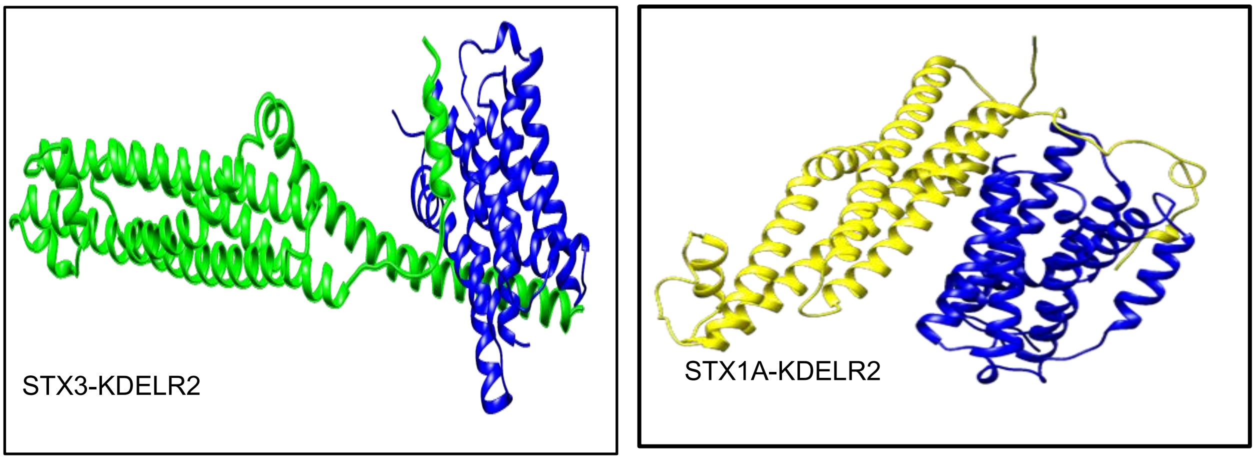


**Sup. Fig.4: 3D structures of STX3 & STX1A with novel interactants after molecular docking**

Interaction of STX3 and STX1A was studied with their novel distinct and common interactants in Haddock and LzerD webservers. The interaction between DNM1 and NSF was taken as positive control.

**Table 4. Computationally predicted amino acid list involved from STX1A and STX3 in the Interaction between novel protein partners**

| No | Interactants Of STXs | Amino Acids involved from STXs in the Interaction Interface | |
|---|---|---|---|
| | | STX3 | STX1A |
| 1 | CORO2B | 54,65,129,136,144,190, 209,210, 211,215, 216, 255, 265, 269 | -- |
| 2 | KIDDIN220 | 114, 185, 186, 223, 244, 260, 261 | -- |
| 3 | CHMP4B | 54, 65,129, 136,144, 190,209, 210, 211, 215, 216, 255, 265, 269 | -- |
| 4 | DNM1 | 57, 65, 125, 133, 221, 222, 223, 256, 257, 281 | 1, 245, 250 , 252, 271, 274, 276 |
| 5 | PKN3 | -- | 233,9,115,29,34,28,38,21,22,23,24,41,20,18,244,243,245,118,122,122,11 |
| 6 | SLC1A3 | -- | 242,27,28,243,32,116,112,30,29,111,248,245,247,246,110,108,103,99 |

| | | | |
|---|---|---|---|
| 7 | NSF | 14,261,260,257,267,268,272,271,274,253,256,17,16,264,275,13 | 170,161,174,172,173,171,177,176,179,168,180,164,160,158 |
| 8 | KRG1 | 264,257,268,271,272,275,279,238,242,259,256,277,281,274,270,273,278 | 236,114,242,237,241,240,238,119,232,2,29,115,111,123,41,37,38,231 |
| 9 | AMBP | 181,282,278,271,267,268,264,263,262,266,270,3,10,6,9,283,279,13,275 | 23, 30 29, 115, 34. 119, 38, 28, 248, 19, 27, 20, 245 |
| 10 | KDELR2 | 256,260,116,10,272,276,275,264,263,267,271,268,3,6,279,9,5,13 | 5,8,6,3,4,2,111,114,42,43,115,34,119,122,41,38,233,243,17,118,126,45,19,29 |

In STX3, amino acids from ~1-28 is NTD, ~29-144 is $H_{abc,}$ ~156-187 is linker domain, and ~194-260 is SNARE motif and in STX1A amino acids from ~30-166 nSec1 binding site, ~158-188 is linker domain, and ~191-270 is SNARE domain.
